# Germline *de novo* mutation rate of the highly heterozygous amphioxus genome

**DOI:** 10.1101/2025.07.14.664012

**Authors:** Jing Xue, Lei Tao, Junwei Cao, Guang Li, Cai Li

**Author notes:** contributed equally (Jing Xue, Lei Tao).

## Abstract

Germline *de novo* mutations (DNMs) are the ultimate source of heritable variation, yet their patterns in highly heterozygous genomes remain poorly understood. Amphioxus, an early-branching chordate with exceptionally high genomic heterozygosity (3.2∼4.2% in sequenced species), offers a unique model to explore mutational dynamics in such contexts. It is unclear whether high heterozygosity in amphioxus is due to a large effective population size, an increased mutation rate, or both. Here, we perform deep short-read whole genome sequencing of a two-generation pedigree of the amphioxus *Branchiostoma floridae* comprising two parents and 104 offspring, and develop a framework based on allele-aware parental assemblies as the reference to accurately identify DNMs. We detect 205 high-confidence DNMs, yielding a genome-wide mutation rate of 5.10 × 10^-9^ per base per generation, which is comparable to that of vertebrates. Combining this estimate with observed nucleotide diversity, we obtain an effective population size of ∼1.9 million, indicating that the elevated heterozygosity mainly results from a large effective population size. We observe a maternal-origin bias when considering all DNMs but a paternal-origin bias for early-occurring ones. Amphioxus harbors a much smaller fraction of CpG>TpG DNMs relative to vertebrates, attributable to its low methylation levels. We also investigate putative post-zygotic mutations in the offspring, revealing an unexpected paternal-origin bias. These suggest some distinct mutational mechanisms in amphioxus. Our study not only provides the first DNM measurement for amphioxus but also offers a generalizable strategy for studying DNMs in highly heterozygous genomes, facilitating mutation rate studies across chordates and other lineages.

## Introduction

Germline *de novo* mutations (DNMs) serve as the ultimate source of genetic variation and are critical for biological evolution. These mutations arise as a consequence of replication errors or mis-repaired/un-repaired DNA damage. The *de novo* mutation rate, the frequency at which DNMs occur, is an important parameter for many genetic and evolutionary analyses. Because most mutations tend to be deleterious, organisms incur considerable adaptive costs to minimize such errors and damage, thereby maintaining mutation rates at an extremely low level, ranging from 10^-11^ to 10^-7^ per base per generation in eukaryotes (Lynch, et al. 2023; Wang and Obbard 2023). Thanks to the advances in sequencing and computational methods, in the past decade, direct detection of DNMs by sequencing parent-offspring trios has been available for many species, particularly for vertebrates (Besenbacher, et al. 2019; Bergeron, et al. 2021; Wang, et al. 2022; Zhang, et al. 2022; Bergeron, et al. 2023; Lin, et al. 2023; Sendell-Price, et al. 2023; Zhang, et al. 2023; Liang, et al. 2024; Porubsky, et al. 2025; Prentout, et al. 2025; Versoza, et al. 2025; Wang, et al. 2025; Zhang, et al. 2025), providing insight into inter-species mutation rate variation and underlying causes.

Mutation rates also vary across genomic positions, developmental stages, parental origins, and mutation types. For example, in mammals the C>T mutation rate at CpG sites is about one order of magnitude higher than the genome-wide average, which is mainly caused by spontaneous deamination of methylated cytosines (Hodgkinson and Eyre-Walker 2011). A recent study in amniotes found that while both sexes exhibit similar mutation rates early in development, the male germline accumulates more mutations after sexual differentiation (de Manuel, et al. 2022), highlighting the influence of gametogenesis on mutation accumulation. Despite the progress, many details of underlying mutational mechanisms remain uncovered.

Amphioxus (cephalochordates) represents the earliest-branching lineage of extant chordates, diverging from the common ancestor of vertebrates and tunicates over 500 million years ago, serving as a key model for understanding vertebrate origins (Holland and Holland 2017). Although amphioxus shares a basic body plan with vertebrates, it is anatomically simpler, and its genome has undergone fewer lineage-specific rearrangements compared to tunicates (Delsuc, et al. 2006; Putnam, et al. 2008). Notably, amphioxus displays an exceptionally high level of genomic heterozygosity (3.2∼4.2% in sequenced species), among the highest observed in chordates (Bi, et al. 2020). Under the neutral theory of molecular evolution, the equilibrium level of heterozygosity is proportional to both mutation rate and effective population size (Kimura 1983). In the absence of accurate mutation rate measurement, it is unclear whether the high heterozygosity in amphioxus is due to a large effective population size, an increased mutation rate, or both.

High heterozygosity is not unique to amphioxus—many species across diverse clades, such as marine invertebrates (Small, et al. 2007; Ketchum, et al. 2020; Nong, et al. 2020; Shingate, et al. 2020; Penaloza, et al. 2021; Wooldridge, et al. 2025), insects (Mackintosh, et al. 2019; Kim, et al. 2021; Bastide, et al. 2022; Garcia-Berro, et al. 2023; Deng, et al. 2024; Jiang, et al. 2025), and plants (Bredeson, et al. 2016; Tang, et al. 2016; Shen, et al. 2018; Wang, et al. 2020; Xian, et al. 2025), also exhibit elevated genomic diversity. Interestingly, multiple sequenced tunicate species such as sea squirts (Small, et al. 2007; Wei, et al. 2020) and larvaceans (Denoeud, et al. 2010; Bliznina, et al. 2021), which are close relatives of vertebrates, also possess genomes with high heterozygosity. Yet, for these species, direct estimates of the germline mutation rate remain unavailable, largely due to technical challenges in working with highly heterozygous genomes (Garimella, et al. 2020). The high heterozygosity presents significant challenges to key genomic analyses such as genome assembly (Kajitani, et al. 2014; Zhang, et al. 2020), read alignment (Landan and Graur 2009), and variant detection (Gan, et al. 2011; Veeckman, et al. 2019). Indeed, previous research in macaques has shown that even gold-standard pipelines for DNM detection exhibit elevated error rates in highly heterozygous genomes (Bergeron, et al. 2022). As a result, the evolutionary forces shaping high genomic diversity in these lineages remain poorly understood.

Amphioxus, with its exceptionally high heterozygosity and relatively small genomes (∼500 Mb) (Huang, et al. 2023), represents an ideal model system for addressing these questions. Direct measurement of DNMs in the amphioxus genome is not only critical for estimating its mutation rate and studying mutational mechanisms, but also highly informative for a broader set of species with extreme genomic diversity.

In this study, we generated deep short-read sequencing data for a two-generation pedigree (two parents and 104 F1 offspring) of *Branchiostoma floridae*, to systematically study DNMs in the amphioxus. To overcome the challenges in alignment and variant detection, we developed a workflow leveraging the high heterozygosity to assemble the contig-level haplotypes of parental genomes, thereby allowing mapping offspring reads to the parental genomes for identifying DNMs. This new and cost-effective strategy can be generalized to other species with high heterozygosity. With the detected DNMs and estimated mutation rate (5.10 × 10^-9^ per base per generation), we concluded that the high heterozygosity in amphioxus is in fact caused by the large effective population size. We further investigated multiple aspects of DNMs as well as postzygotic mutations (PZMs) in amphioxus, providing insights into mutational mechanisms in this basal chordate lineage.

## Results

### An analysis strategy based on parental genome assembly for detecting DNMs

To identify DNMs, we performed deep short-read whole genome sequencing (WGS) for two parental individuals and 104 F1 offspring of a *B. floridae* family, yielding an average coverage of 83× per sample (ranging from 49× to 141×) (**Supplementary Figure 1 and Supplementary Table 1**).

*k*-mer analysis revealed that the heterozygosity of the parental genomes was approximately 3% (**Supplementary Figure 2**). The high level of heterozygosity in the *B. floridae* genome poses a challenge for conventional DNM detection pipelines based on aligning reads of parents and offspring against a public reference genome (Bergeron, et al. 2022). Because of low sequence similarity between our sequencing data and the public reference genome (Huang, et al. 2023), many reads in our sequencing data cannot be correctly mapped to the reference genome by common short-read aligners (e.g. BWA), leading to unreliable variant calling results. By simulating sequencing reads with ART (Huang, et al. 2012), we estimated that, when the sequence difference between the query genome and the reference genome is 4%, >2% of reads are mapped to wrong genomic regions (**Supplementary Table 2**) and such problematic alignments are difficult to distinguish in real datasets when calling variants.

However, we reasoned that the high heterozygosity of *B. floridae* can facilitate allele-aware assembly of the parental genomes, as many divergent regions in the parental genomes can be assembled into separated contigs. The high sequencing depths of our data enabled us to do contig-level *de novo* assembly for two parents. Using the parental genome assembly as a custom reference, we should be able to align most offspring and parental reads to the parental genomes, thereby obtaining high-quality genotype information of offspring and parents for DNM detection (**Figure 1**).

**Figure 1.**
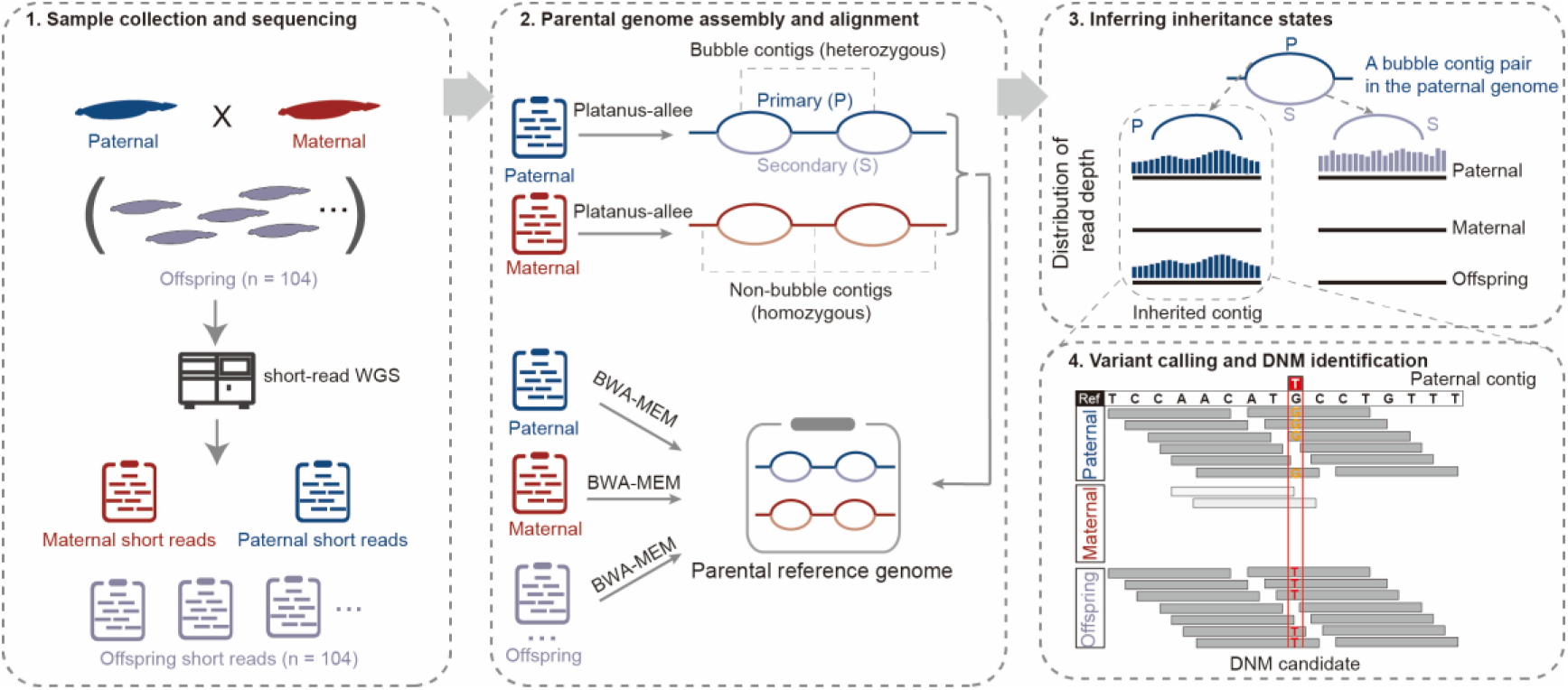
Schematic of the workflow for detecting DNMs. We collected DNA samples from a *B. floridae* family of two parents and 104 F1 offspring and performed deep short-read sequencing (>50×) for all individuals. Using the parental WGS data, we assembled allele-aware parental genomes with Platanus-allee (Kajitani, et al. 2019) and combined the resulting assemblies to form a parental reference genome for downstream analyses. Short reads from both parents and offspring were aligned to this custom reference using BWA-MEM2 (Vasimuddin, et al. 2019). We then computed the read depth distribution for each contig in both parents and offspring. An example is shown for a pair of paternal bubble contigs. Based on the read depth distributions in parents and an offspring, we identified the inherited contig from the father in the offspring. The inheritance states of all parental contigs and read alignments were further used to identify DNMs in each offspring, with an exemplar DNM shown in the bottom right panel.

Therefore, we constructed the genome assembly for each parent using the allele-aware assembler Platanus-allee (v2.2.2) (Kajitani, et al. 2019). We obtained a maternal genome assembly of 733.8 Mb and a paternal genome assembly of 763.7 Mb (**Supplementary Table 3**), approximately 1.6 times the length of the previously published haploid genome (490 Mb) (Huang, et al. 2023), indicating that a large fraction of the diploid parental genome was assembled into phased alleles. By the definition of Platanus-allee, heterozygous regions are assembled as “bubble” contigs with two allelic sequences, named as “primary” and “secondary” respectively. Homozygous regions with identical or nearly identical sequences in an individual are assembled as “non-bubble” contigs (**Figure 1**).

By merging the two parental genome assemblies, we generated a composite parental reference genome, with detailed assembly statistics provided in **Supplementary Table 3**. The parental reference genome had a total length of approximately 1.4 Gb, consisting of 263,358 bubble contigs covering ∼856 Mb and 758,976 non-bubble contigs spanning ∼780 Mb. The contig N50 metrics of maternal and paternal assemblies were 6.6 kb and 7.6 kb, respectively. >91% of sequencing reads of the two parents were aligned to this parental reference genome with proper insert sizes (**Supplementary Figure 1**), indicating that the vast majority of genomic segments were successfully assembled and available for downstream analysis.

Next, we aligned sequencing reads from both offspring and parents to this parental reference genome and detected variants using BCFtools (v1.21) (Danecek, et al. 2021) and freebayes (v1.3.10) (Garrison and Marth 2012). A custom filtering pipeline, accounting for factors such as read depth and variant quality, was employed to obtain high-confidence DNMs (see Methods). After applying the filtering criteria, 321 candidate DNMs were retained. Among these DNMs, putative false positives were further ruled out by manual check with IGV visualization (see Methods). After careful check, a total of 205 high-confidence DNMs (**Supplementary Table 4**) were retained across 104 offspring, corresponding to an average of 1.97 DNMs per individual. 22 DNMs from 9 individuals were selected for orthogonal validation using independent amplicon-seq data. For each selected DNM, 150∼200 bp regions encompassing the DNM were amplified and sequenced for both the offspring and parents. The amplicon-seq data confirmed 20 of the 22 selected DNMs as true DNMs (**Supplementary Table 5-7**), while the remaining two (in a same amplicon) yielded inconclusive results due to amplification noises, indicating a low false positive rate in our DNMs. With an average callable genome size of 398.6 Mb and a false negative rate of 3% estimated by simulation (Methods), the mean genome-wide mutation rate was calculated to be 5.10×10^-9^ mutations per base per generation, with a 95% confidence interval (CI) of 4.46-5.83×10^-9^. The mutation rate of *B. floridae* is comparable to those reported in fish and other vertebrates (Bergeron, et al. 2023) (**Figure 2A**).

**Figure 2.**
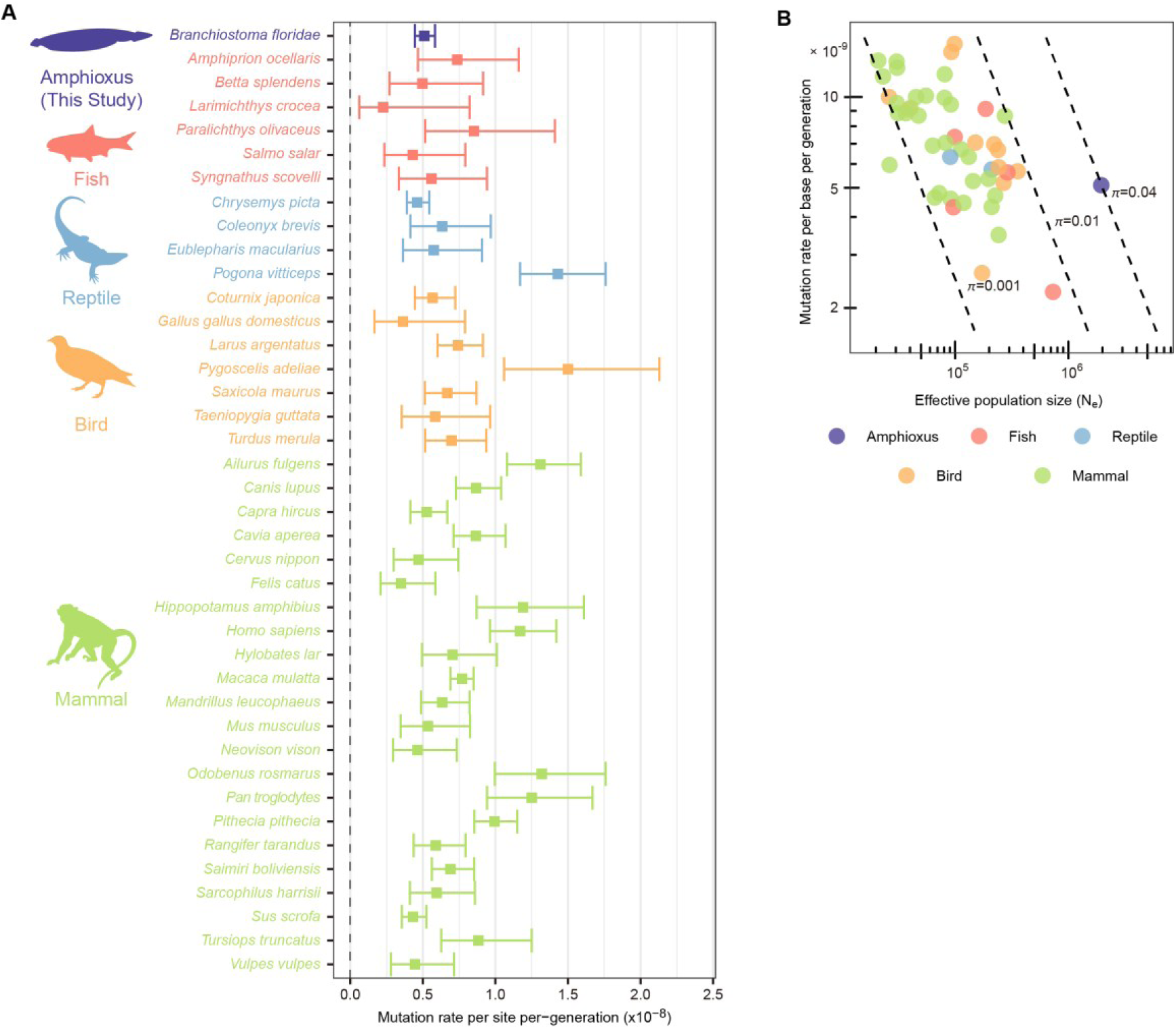
Estimates of mutation rate and effective population size. (**A**) Per base per generation *de novo* mutation rate (μ) estimates from this study and previous studies (Bergeron, et al. 2023). Error bars denote 95% confidence intervals. (**B**) Comparison of mutation rates and effective population sizes across diverse taxonomic groups, using data for other taxa from Bergeron et al. (2023). The dashed lines from left to right represent π values of 0.001, 0.01, and 0.04, respectively. The π values, labeled near each dashed line, were calculated according to π = 4N_e_μ.

### High heterozygosity of *B. floridae* is mainly attributed to its large effective population size

It has long been recognized that the genomic heterozygosity of *B. floridae* is much higher than in other species (Putnam, et al. 2008). Yet, since the mutation rate in *B. floridae* has not been directly quantified, the cause(s) of its high heterozygosity remains uncertain. The nucleotide diversity, expressed as π, can be represented by π = 4N_e_μ under neutral evolution (Nei and Tajima 1981; Kimura 1983), where N_e_ represents the effective population size and μ the germline mutation rate. Therefore, the effective population size can be estimated by the equation N_e_ = π/(4μ). Based on synonymous substitutions in the population genomic data from Huang et al. (2023), we obtained the π estimate of 0.0397 for *B. floridae* (Methods). For comparison, the π estimates of most vertebrate species from Bergeron et al. (2023) fall into the range of 0.001 to 0.01, with none above 0.01. The effective population size, calculated with our estimated mutation rate (μ) and π, was 1,946,770. We placed *B. floridae*’s N_e_ and μ within the context of N_e_ and μ values derived from Bergeron et al. (Bergeron, et al. 2023) to show its unique position (**Figure 2B; Supplementary Table 8**), indicating that *B. floridae*’s N_e_ is largest among chordates with measurements. We note that, the sequenced population in Huang et al. (2023) experienced some degree of inbreeding in the laboratory, which could lead to lower nucleotide diversity than wild populations. Therefore, the actual N_e_ in wild populations is likely larger than the current estimate. Collectively, we concluded that the high heterozygosity of amphioxus is not due to an elevated mutation rate but a very large effective population size.

### Characteristics of DNMs in *B. floridae*

Distinguishing the parental origins of DNMs is crucial for understanding mutational dynamics. While previous studies could only assign parental origin for 30%∼80% of DNMs based on nearby polymorphic markers (Bergeron, et al. 2023; Wooldridge, et al. 2025), our DNM calling method inherently provided parental origin information for all detected DNMs. Among the 205 DNMs, 105 originated from the mother and 100 from the father (**Figure 3A**), yielding a male mutation bias coefficient (α) of 0.95. Considering the difference in the callable genome sizes between the parents, we obtained a corrected α of 0.88 (95% CI = 0.83-0.91), indicating a maternal bias. For a cross-species comparison, we integrated several representative vertebrates with known α values (**Figure 3B**). The results showed that the α value of *B. floridae* is lower than most vertebrates. Previous studies (Bergeron, et al. 2023; Zhang, et al. 2025) reported that fish tend to have apparently lower α values than other vertebrates (mammals, birds and reptiles). These studies suggested that differences in gametogenesis such as seasonal breeding and synchronous ovulation could lead to the lower α values in fish. The low α of amphioxus might be partly due to these reasons.

**Figure 3.**
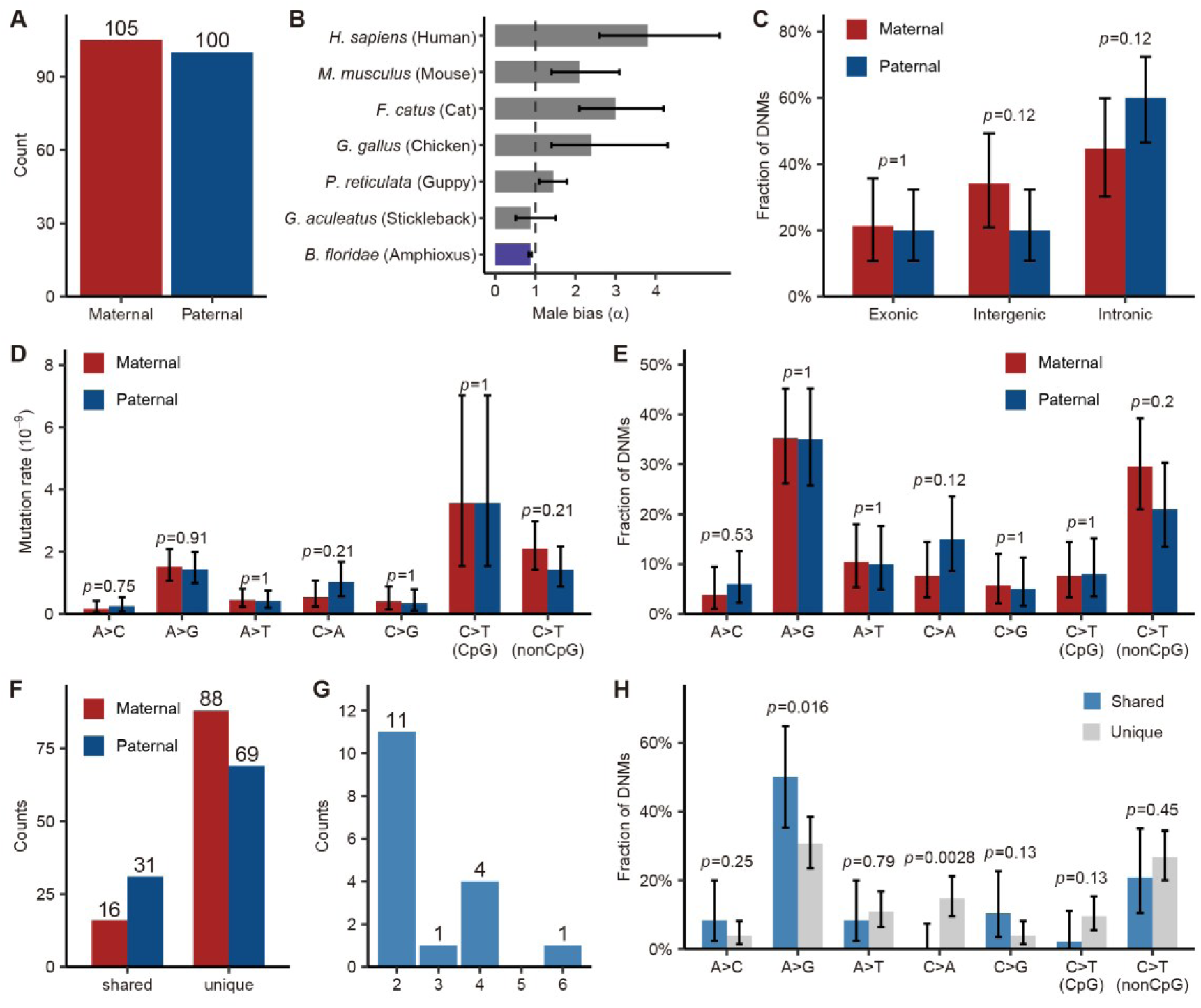
Characteristics of DNMs in *B. floridae*. **(A)** Parental origins of DNMs. **(B)** The male-to-female ratios (α) of DNMs for different chordates. **(C)** Proportions of DNMs in different gene annotation contexts, separating paternal (blue) and maternal (red) DNMs. Parental origin differences were assessed by Fisher’s exact tests. **(D)** Mutation rates of different substitution types, separating paternal (blue) and maternal (red) DNMs. Parental origin differences assessed by χ² tests. The mutation spectrum was folded (e.g., A>C representing both A>C and T>G mutations). **(E)** Proportions of DNMs of different substitution types, separating paternal (blue) and maternal (red) DNMs. Parental origin differences were assessed by Fisher’s exact tests. **(F)** Counts of sibling-shared and unique DNMs, separating paternal (blue) and maternal (red) DNMs. **(G)** Counts of sibling-shared DNMs shared by different numbers of offspring. **(H)** Proportions of DNMs of different substitution types, separating shared and unique DNMs. Differences between shared and unique DNMs were assessed by Fisher’s exact tests. Error bars denote 95% confidence intervals computed using binomial distributions. All p-values for Fisher’s exact tests and χ² tests are given above the bars.

Due to the fragmented nature of the parental genome assemblies, direct genomic annotation was challenging; therefore, we mapped the parental genome coordinates to the published haploid reference genome from Huang et al. (Huang, et al. 2023) and used its annotation for downstream functional analyses. Of the total 490 Mb of the public haploid reference genome, 363 Mb (74%) was covered by the assembled contigs of parental genomes. The parental contigs that were mapped to the haploid reference genome were defined as mappable regions with annotation information. Based on the haploid genome annotation (Huang, et al. 2023), the genome was divided into three gene annotation contexts: exonic, intronic, and intergenic regions. We found no significant difference between the fractions of paternal and maternal DNMs for each gene annotation context (**Figure 3C**).

We further investigated DNM counts of different substitution types. The results indicated no significant difference between maternal and paternal mutation rates for all substitution types (**Figure 3D**), and no significant difference between parental origins when looking at proportions of DNMs of different substitution types (**Figure 3E**). In vertebrates, the mutation rate of C>T at CpG sites is usually much higher than that at non-CpG sites (e.g., >10 fold difference in humans) (Kong, et al. 2012; Francioli, et al. 2015). However, the mutation rate of C>T at CpG sites was only about twice that at non-CpG sites in *B. floridae* (**Figure 3D**). Cross-species comparison of mutational spectra (**Supplementary Figure 3**) revealed some unique patterns in *B. floridae*. CpG>TpG mutations only accounted for 8% of DNMs in *B. floridae*, significantly lower than those (16-20%) observed in vertebrates. On the other hand, A>G mutations accounted for 35% of DNMs in *B. floridae*, consistently higher than those (16-26%) in surveyed vertebrates. The lower mutation frequency at CpG sites was probably due to lower methylation levels at CpG sites in amphioxus (Marletaz, et al. 2018).

47 (23%) DNMs were shared among siblings (named ssDNMs, **Figure 3F**), which likely occurred at early developmental stages. A greater number of ssDNMs were identified as paternal in origin, with 31 ssDNMs derived from the father compared to only 16 from the mother. The male bias α of 1.80 (after accounting for callable size difference), was substantially higher than the α of 0.88 observed for all DNMs (Fisher’s exact test, p = 0.036), suggesting that during earlier stages of germ cell development, male germline accumulates more mutations relative to female germline. Majority of ssDNMs were shared by only two offspring, with one shared by up to six offspring (**Figure 3G**). Comparison of mutation spectra indicated a significant increase in the proportion of A>G mutations in ssDNMs relative to unique DNMs, but a depletion of C>A mutations in ssDNMs, likely reflecting the differences in mutational processes between early and late gametogenesis (**Figure 3H**).

### Characteristics of PZMs in *B. floridae*

Postzygotic mutations (PZMs) arise during mitotic cell divisions in the developing embryo or later in life, and thus typically affect only a subset of the body’s cells. Some PZMs are present in the germ cells can be passed to the next generation. The high sequencing depths of our data also enabled us to detect PZMs in offspring (see Methods for details). We identified 204 putative PZMs across 104 offspring (**Figure 4A; Supplementary Table 9**), yielding an average of 1.96 PZMs per individual. The variant allele frequencies of PZMs ranged from 0.05 to 0.45 (**Supplementary Table 9**). As the detected PZMs had relatively high allele frequencies, they probably arose during early development.

**Figure 4.**
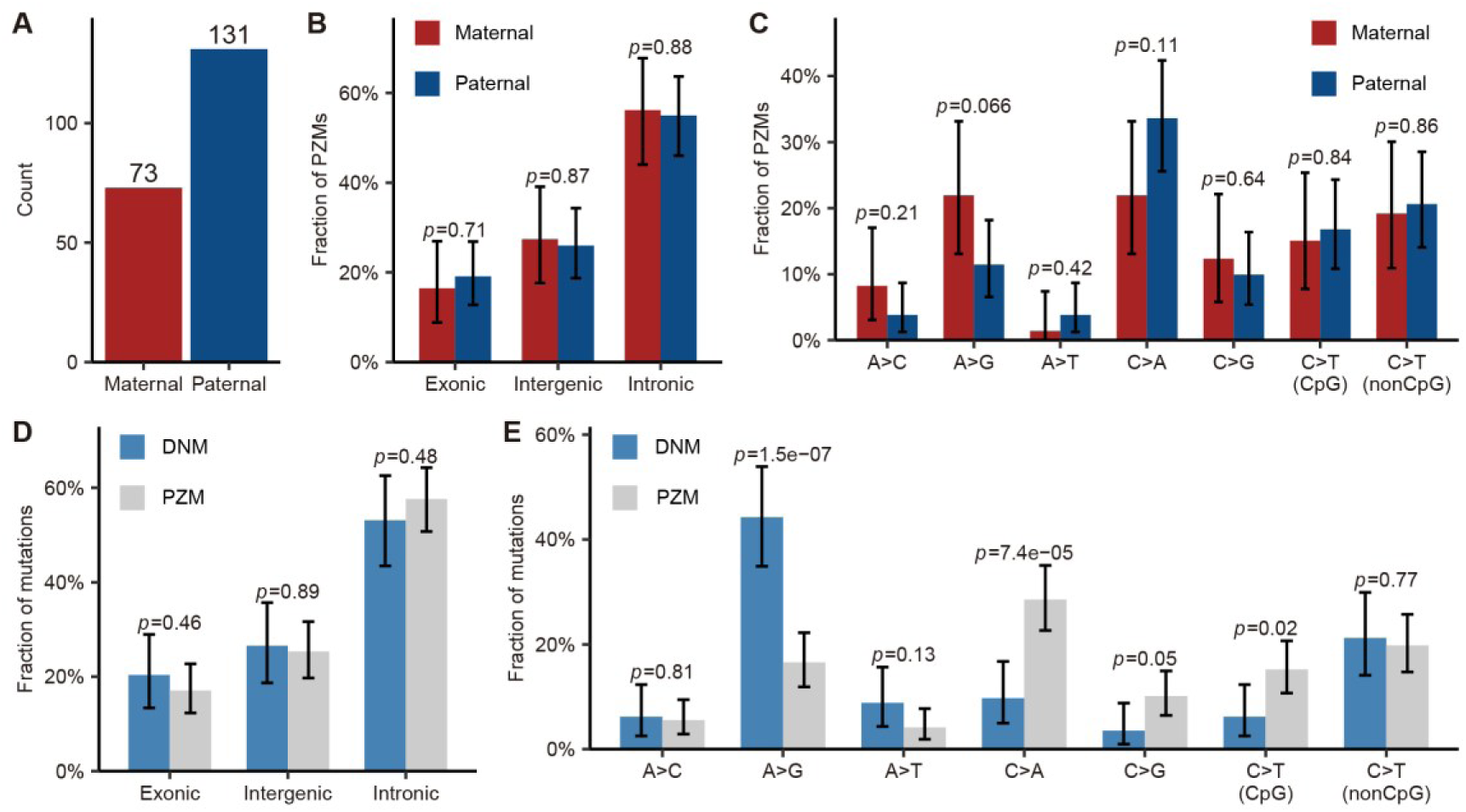
Characteristics of PZMs. (**A**) Parental origins of PZMs. (**B**) Proportions of PZMs in different gene annotation contexts, separating paternal (blue) and maternal (red) PZMs. Parental origin differences were assessed by Fisher’s exact tests. (**C**) Proportions of PZMs of different substitution types, separating paternal (blue) and maternal (red) PZMs. (**D**) Proportions of mutations in different gene annotation contexts, separating DNMs (blue) and PZMs (grey). **(E**) Proportions of mutations of different substitution types, separating DNMs (blue) and PZMs (grey). Parental origin differences and differences between DNMs and PZMs were assessed by Fisher’s exact tests, with p-values above the bars. Error bars denote 95% confidence intervals computed based on binomial distributions.

Unexpectedly, PZMs in *B. floridae* exhibited a pronounced paternal origin bias (α = 1.67, corrected for callable size difference; binomial test, p = 6.9e-07; **Figure 4A**). The sex-biased pattern of PZMs, which has not been observed in other species to our knowledge, suggested different mutational processes operating on paternal and maternal chromosomes during early development. In mammals, early divisions of embryos erase epigenetic memory (primarily DNA methylation and histone modifications) from sperm and egg and re-establish the epigenomic environment. Yet in some vertebrates such as zebrafish and sea lamprey (Potok, et al. 2013; Angeloni, et al. 2024), the methylation landscape in the paternal genome of early embryos is largely maintained and the maternal methylation is reprogrammed to match the paternal. Available methylome data suggested that similar patterns are likely to be present in sea squirt and sea urchin (Xu, et al. 2019). While there was no study comparing methylomes of gametes and early embryos in amphioxus, we suspected that amphioxus may employ similar epigenetic reprograming like that in sea lamprey, given its phylogenetic position. The epigenomic differences between paternal and maternal chromosomes in early embryos might contribute to different frequencies of paternal and maternal PZMs.

We further examined differences in parental origin with respect to proportions of PZMs across gene annotation contexts and across mutation types, but found no significant difference (**Figure 4B-C**). Similarly, comparison between PZMs and DNMs in gene annotation contexts revealed no notable difference (**Figure 4D**). However, analysis of specific mutation types revealed that A>G mutations were significantly more frequent in DNMs than in PZMs. Conversely, C>A mutations were markedly enriched in PZMs relative to DNMs, and C>G and CpG>TpG mutations were also more prevalent in PZMs (**Figure 4E**). Previous methylome analysis revealed that amphioxus embryos exhibit more methylated CpGs relative to adult tissues, which may partly explain the relatively higher frequency of CpG>TpG mutations in PZMs (Marletaz, et al. 2018). While the parental origin ratio of PZMs was close to that of ssDNMs (PZM: 1.67; ssDNM: 1.80), their mutation spectra were largely different (**Figure 3H** versus **Figure 4E**), suggesting different underlying mutational mechanisms.

## Discussion

In this study, we generated high-depth short-read sequencing data for a pedigree comprising both parents and 104 offspring to investigate DNMs in amphioxus. Due to high heterozygosity of amphioxus, which results in many sequence differences between the public reference genome and re-sequenced individuals, conventional alignment and variant calling methods that rely on the public reference genome are prone to errors, thereby complicating DNM detection. However, high heterozygosity can also facilitate allele-aware diploid genome assembly without requiring long-read data. With this consideration, we assembled haplotype-resolved diploid genomes at the contig level for each parent and developed a custom read mapping and variant calling pipeline using the parental assemblies as the reference. This cost-effective strategy can help resolve the challenges of detecting DNMs in highly heterozygous species like *B. floridae*. It also allowed us to assign the parent-of-origin state for all detected DNMs. Furthermore, based on phased genotypes of parents and offspring, we were also able to comprehensively investigate the meiotic recombination landscape, which has been described in a companion study (Tao, et al. 2025).

With an average callable diploid genome size of 398.6 Mb, we identified 205 DNMs and estimated the germline mutation rate in *B. floridae* to be 5.10 × 10^-9^ (95% CI of 4.46-5.83 × 10^-9^) per base per generation. Based on this mutation rate, the effective population size of *B. floridae* was estimated to be about two million, suggesting that the high heterozygosity resulted from a massive effective population size. Our estimated mutation rate for contemporary populations is close to a recently reported estimate (4.65 × 10^-9^) based on divergence times and intra-cephalochordate synonymous substitution rates of single-copy genes (Ren, et al. 2025). Our study did not estimate the germline mutation rate for small insertions/deletions (INDELs), as generally poor alignments around these variants (usually associated with repeats) make it challenging to accurately detect *de novo* INDELs. In addition, repetitive or low-complexity regions, which usually have relatively high mutation rates, are difficult to assemble with only short reads and thus were under-represented in the callable regions of our study. Future studies incorporating long-read sequencing technologies (e.g., PacBio or Oxford Nanopore) can facilitate identifying *de novo* INDELs as well as other DNMs in complex genomic regions.

We further analyzed the parental origin, genomic distribution, mutation spectrum, and ssDNMs in *B. floridae*, revealing some distinctive features in this basal chordate. We observed a maternal bias of DNMs, with a male-to-female mutation ratio (α) of 0.88. This ratio was lower than most reported values for vertebrates, likely associated with seasonal breeding and synchronous ovulation (de Manuel, et al. 2022; Bergeron, et al. 2023). The overall mutation spectra showed no obvious difference between different parental origins. The low frequency of C>T transitions at CpG sites was consistent with previously reported low levels of CpG methylation in amphioxus (Marletaz, et al. 2018). The presence of ssDNMs provided a valuable window into early germline mutational events. We observed more ssDNMs in the paternal genome relative to the maternal, but the underlying cause is unclear. Future DNM studies spanning different ages and developmental stages of parents can help clarify the mechanisms of mutagenesis in amphioxus. In addition, functional genomics studies focusing on DNA damage and repair pathways are also essential to understanding the mutational mechanisms.

We identified 204 putativePZMs across 104 offspring, with a surprisingly strong paternal-origin bias (α = 1.67). As the bias was unlikely due to artifacts after careful check, we speculated that the paternal bias of PZMs is likely linked to underlying epigenomic asymmetries between parental genomes in early embryos—such as incomplete or asymmetric reprogramming of DNA methylation—which may lead to different susceptibility to mutations. Further functional genomic analysis is needed to test this hypothesis. While PZMs and DNMs showed similar distributions across gene annotation contexts, PZMs were significantly enriched for C>A and CpG>TpG mutations, whereas germline DNMs more frequently involved A>G mutations. As most PZMs are somatic mutations, the distinct mutation spectra between PZMs and DNMs highlighted different biological mechanisms governing mutagenesis in somatic versus germline lineages.

In summary, through high-depth pedigree sequencing and the development of a novel analysis pipeline, we directly quantified the germline mutation rate in *B. floridae* for the first time and resolved the long-standing question of what causes its high heterozygosity. We also explored DNMs from multiple perspectives, including parental origin, gene annotation context, mutation spectrum and shared DNMs among siblings. Some of the unique DNM features observed in amphioxus may represent ancestral characteristics of germline mutagenesis in chordates. The analysis framework based on assembling parental genomes for read alignment and variant calling can facilitate related research in other highly heterozygous species.

## Materials and Methods

### Amphioxus culture

Amphioxus (*B. floridae*) strains obtained from a stock maintained by Dr. Jr-Kai Yu originating from Tampa, Florida, were housed in a temperature-controlled facility at Xiamen University, which maintained a 14 h light and 10 h dark cycle and ambient seawater temperature at 19°C (Li, et al. 2012). A pair of parental individuals was selected for crossing. Fertilized eggs were subsequently cultured following the cultivation strategy described previously until they reached the designated sampling stage. A total of 104 F1 offspring were generated through this process. To capture distinct developmental time points, the offspring were snap-frozen in liquid nitrogen at two specific post-growth intervals: 6 months and 1 year. The first batch, comprising 35 individuals, was preserved at the 6-month time point, and the second batch, consisting of 71 individuals, was preserved at the 1-year time point.

### Library construction and sequencing

We extracted high molecular weight genomic DNAs from the muscle tissues of a single individual using the DNeasy Blood & Tissue Kit (QIAGEN, Valencia, CA), and inspected the DNA quality by Qubit 2.0 Fluorometer (Thermo Fisher Scientific, Waltham, MA) and 2100 Agilent Bioanalyzer (Agilent). After sample QC, gDNA was fragmented by ultrasound on Covaris E220 (Covaris, Brighton, UK), then selected to 300-500 bp using magnetic beads size selection. The selected DNA fragments were then end-repaired and A-tailed. The indexed adaptors were ligated to both ends of the DNA fragments. The ligation product was amplified by PCR, and circularized to get single-stranded circular (ssCir) library. The ssCir library was then amplified through rolling circle amplification (RCA) to obtain DNA nanoball (DNB). The DNB was then loaded to flowcell, and sequenced by DNBSEQ Platform (MGI, China).

### Parental genome assembly

Genome heterozygosity of parental individuals was estimated via *k*-mer analysis of Illumina short-read data using GenomeScope (v2.0) (Ranallo-Benavidez, et al. 2020). Raw short-read sequences were quality-processed with fastp (v0.24.0) (Chen 2023), implementing: (1) elimination of reads with >10% ambiguous bases (’N’), and (2) retention of reads exhibiting a minimum mean Phred quality score of 20 (Q20). Adapter sequences were concurrently trimmed. These processed reads were subjected to allele-aware diploid genome assembly using Platanus-allee (v2.2.2) (Kajitani, et al. 2019) through a two-stage workflow: First, the *de novo* assembly phase was executed with the command “assemble -m 200 -t 20”; second, haplotype resolution was performed via the phase module using “phase -i 3 -t 20”. Contigs shorter than 150 bp in length were subsequently filtered from both parental phased assemblies. Initial assembly quality metrics, including contig N50, total length, and GC content, were comprehensively evaluated with QUAST (v5.3.0) (Gurevich, et al. 2013) and SeqKit (v2.9.0) (Shen, et al. 2016).

Following the phased assembly procedure, contigs underwent automated classification by Platanus-allee. In the contig-level assemblies generated by Platanus-allee, contigs were classified into two distinct categories: (1) bubble contigs harboring allelic variants, which were further stratified as primary contigs (longer allelic sequences) and secondary contigs (shorter counterparts), and (2) non-bubble contigs designated by the assembler as lacking allelic diversity. However, through comprehensive whole-genome alignment analyses, we discovered that a subset of non-bubble contigs exhibited unexpected sequence overlaps with other contigs across the assembly, suggesting residual heterozygous regions unresolved by the phasing algorithm.

Finally, the filtered phased diploid contig sets derived from paternal and maternal genomes were concatenated to construct a custom reference genome (hereafter termed parental reference genome), which were used in many downstream analyses.

### Alignment of offspring and parental reads to the parental reference genome and variant calling

Processed short reads from two parents and offspring were aligned to the parental reference genome assembly using BWA-MEM2 (v2.2.1) (Vasimuddin, et al. 2019) with parameters “-M -k 19”. Post-alignment processing entailed coordinate-based sorting and removal of PCR duplicates via SAMtools (v1.21) (Li, et al. 2009) and Picard Tools in GATK4 (v4.6.1.0) (Van der Auwera and O’Connor 2020) with default parameters. Per-base depths across the whole reference genome and individual-level average sequencing depth metrics were subsequently computed using mosdepth (v0.3.10) (Pedersen and Quinlan 2018) with default parameters.

Based on the read alignments above, we used BCFtools (v1.21) (Danecek, et al. 2021) and freebayes (v1.3.10) (Garrison and Marth 2012) to identify small variants. First, we generated “pileup” files for the set of all samples by running “bcftools mpileup” with parameters “-annotate FORMAT/AD, FORMAT/ADF, FORMAT/ADR, FORMAT/DP, FORMAT/SP, INFO/AD, INFO/ADF, INFO/ADR” on all input bam files. With the “pileup” files as input, we ran “bcftools call -mv” to generate VCF files, which contain SNVs and short insertions/deletions, for 104 offspring and two parents. We additionally used freebayes (v1.3.10) (Garrison and Marth 2012) to identify variants within similar sequences between the two parents with parameters “-m 0 -- legacy-gls.” In regions exhibiting high sequence similarity between the two parents, only the freebayes calls were retained; in all other regions, variant calls from both tools were used.

### Inferring inheritance states of parental contigs in offspring genomes based on read alignments against the parental reference genome

According to the principles of Mendelian inheritance, genomic reads in the offspring are derived from the gene pool of two parental genomes. As described in previous sections, we obtained the paternal and maternal diploid genome assemblies as sets of haplotype-resolved contigs. By mapping sequencing reads of an offspring back to the parental reference genome (combining paternal and maternal genome assemblies), we can determine which allele of a specific locus (bubble or non-bubble contigs) of a given parent is inherited by the offspring. More specifically, to determine whether a particular contig from a given parent is inherited by a specific offspring, we compared read depth distribution profiles of that contig between the offspring and each parent. For a contig with specific parental origin, the depth distribution of the offspring’s reads aligned to this contig is expected to more closely resemble the depth distribution of the reads from the parent-of-origin rather than that from the opposite parent.

The workflow for inferring inheritance states is described below and illustrated in **Supplementary Figure 4**. To quantify the depth distribution along a contig, each contig was divided into 50 bp bins, discarding any terminal bin shorter than 50 bp. The mean depth within each bin was calculated for each individual, generating a depth distribution vector per individual across the contig. The similarity between offspring and parental vectors was quantified using a weighted measure that combines (1) Cosine similarity for bins with non-zero depth in both individuals being compared, and (2) Jaccard similarity for bins with zero coverage in at least one of the two individuals (accounting for regions of no aligned reads). Weights of different similarity scores were allocated based on corresponding proportions of zero-value bins and non-zero-value bins. A parental contig was considered inherited by an offspring if its depth distribution vector in that offspring showed greater similarity to the vector of its specific parental source than to the other parent. Parental contigs exhibiting a depth less than 1/4 of an offspring’s mean depth or a depth less than 10 were classified as low-depth and thus non-inherited in the specific offspring.

However, the similarity of some contigs between the parental genomes, probably arising from shared ancestry, could confound the inheritance calls derived from the offspring-parent depth distribution similarity algorithm (**Supplementary Figure 5**). Since direct pairwise genome sequence alignment of two parental genomes is difficult for assessing parental contig similarities due to potential length disparities of orthologous contigs in different parental assemblies, we used the similarity of read depth distributions (calculated using the same algorithm employed for offspring-parent comparisons) when mapping each parent’s reads to the same contig as a robust proxy. If the depth distribution similarity between the two parents for a specific contig is greater than 0.5, the inheritance calls for that contig in the offspring are deemed unreliable due to indistinguishable parental origins, necessitating further validation with additional data.

Because a given diploid offspring can inherit only one allele from each parent at a locus, determining the inheritance status of one contig within a pair of bubble contigs from a parent inherently resolves the inheritance status of both contigs in the pair. In other words, the inheritance of one contig in a bubble pair can be inferred based on the inheritance of its counterpart. Therefore, we further took advantage of the very low read depths in some loci to conclude that an allele was not inherited; under such circumstances, the alternative allele with aligned reads must be inherited by the offspring. In contrast, for non-bubble contigs, the inability to identify a second allele precludes this method of assessing similarity between parental contigs, and therefore these contigs were excluded from further analysis. Finally, all scenarios arising from the combinations of inheritance and contig similarity, which lead to differences in depth distributions in offspring, were comprehensively illustrated in **Supplementary Figure 5**.

Based on the inheritance states of parental contigs in each offspring, we in fact have reconstructed the offspring genomes with phased alleles in the majority of regions. Such phased offspring genomes can facilitate downstream analyses.

### Determining the callable genome

Estimating mutation rates requires determining the length of genomic regions within each offspring in which DNMs can be confidently identified, defined as the callable genome size. In each offspring, the callable genome is primarily composed of the inherited genomic segments defined in the previous section. However, the accurate inference of the inheritance of non-bubble contigs with high similarity between the two parents proved challenging. For non-bubble contigs lacking pre-existing allelic data in the parental genomes, the allele-depth-distribution-based method for inferring contig inheritance patterns is inapplicable. Consequently, determining the inheritance of homologous contigs within non-bubble contigs becomes challenging. We therefore implemented stricter filtering criteria, excluding non-bubble contigs with parental depth distribution similarity ≥0.5 from the callable genome regions.

Regions with high sequence similarity between parental alleles impede the variant detection and phasing in our pipeline. In this study, the parents were derived from an inbred population, and thus their offspring could inherit highly similar or identical alleles from parents in some regions. To address this, we developed a pipeline to identify pairs of parental contigs with high sequence similarity. Briefly, we performed pairwise genome alignments using Minimap2 (v2.24) (Li 2021) with parameters “-cx asm20 --secondary=no”, designating the paternal genome as the reference and the maternal genome as the query, and *vice versa*. Reciprocal best-hit contig pairs were taken as evidence of homology. For each pair, we further assessed the depth distribution similarity based on alignments of parental reads to assembled contigs; a pair of contigs with a similarity score >0.5 was classified as a highly similar allele pair. For each offspring, if both contigs of a highly similar allele pair were inherited by the offspring, they were excluded from the callable genome for that offspring, as DNMs in these contigs could not be reliably phased and could be confounded by PZMs. In contrast, when only one homologous contig from either parent of a highly similar allele pair was inherited, the contig was retained in the callable genome with the freebayes-called variants.

Additionally, we removed sites in repetitive regions, as they are prone to erroneous alignments and often exhibit abnormally high alignment depths. Sites with too many reads may represent problematic repetitive regions (Li 2014). Specifically, based on the observed depth distribution in each offspring, any regions with a read depth exceeding three times the individual’s average were excluded. The terminal regions of contigs (100 bp at each end) were also excluded from the callable genome, because these regions tended to enrich assembly and alignment errors.

Finally, by integrating these filters based on both inheritance and sequencing depth, we obtained the callable genome size for each offspring ranging from 334.11 Mb to 463.06 Mb, accounting for approximately 34-47% of the diploid genome size (**Supplementary Table 10**).

### Identifying the candidate DNMs

For each parent-offspring trio, the variants in three VCF files of each trio were filtered to a subset of single nucleotide variants (SNVs) by BCFtools (v1.21) (Danecek, et al. 2021) and freebayes (v1.3.10) (Garrison and Marth 2012).

Variants were retained only if they met all of the following criteria: (1) the variant was located within the predefined callable genome regions; (2) the variant had homozygous alternative allele status in the offspring’s alignment against the contig with read depth >10, and read depth of the inherited parent at the variant site of the contig was required to be >5; (3) there were at most two alternative alleles at the variant site in each parental BAM file; (4) the variant had a minimum QUAL score of 220, ensuring high-confidence base calls (BCFtools only); (5) no proximity to indels (outside ±20 bp of indels); (6) variants not at the sites experiencing putative gene conversions described in a companion study (Tao, et al. 2025), as gene conversions lead to read alignments resembling DNMs; (7) for a candidate DNM detected on a specific parental contig in an offspring, all other offspring did not contain a heterozygous genotype at the same site of the contig, as heterozygosity implies an inherited variant or alignment errors.

After applying these filters, the resulting set of candidate DNMs was further evaluated by importing both parental and offspring BAM files into IGV (v2.19.2) (Thorvaldsdottir, et al. 2013), and using IGVtools to generate screenshots for each candidate DNM in both the offspring and the parents. Subsequent manual inspection led to the removal of any candidate DNMs if they met one or more of the following criteria: (1) structural variant/recombination breakpoint neighborhoods, and (2) variants were found to arise from misassigned parental origin of contigs, upon manual review. These steps yielded a DNM dataset for each individual (**Supplementary Table 4**). The genome-wide mutation rate was calculated by the number of all DNMs divided by the sum of callable genome sizes of all offspring. The callable genome size was not multiplied by two, since the reconstructed inherited genomic segments effectively represent an approximately diploid genome for each offspring.

Because the coordinates of DNMs were located on the parental genomic contigs, the parental origin of the contig harboring each DNM naturally revealed its source, allowing for unequivocal determination of the DNM’s parental origin without any further processing. This same principle applied to the callable genome as well.

### Simulation for assessing alignment errors if using the public reference genome

To assess alignment errors due to low sequence similarity between our sequencing data and the public *B. floridae* reference genome, we performed a simulation analysis. Artificial genomes were generated by introducing varying proportions of substitutions into the published *B. floridae* reference. Paired-end 150 bp reads were then simulated from these artificial genomes using ART (Huang, et al. 2012) with parameters “-p -m 450 -s 50 -f 10 -l 150 -ef -ss HS25”. The simulated reads were aligned to the original *B. floridae* reference using BWA-MEM2 (v2.2.1) (Vasimuddin, et al. 2019), and mis-mapped reads were quantified with a custom script. The results were summarized in **Supplementary Table 2**.

### Simulation for assessing the false negative rate of DNM detection

For evaluating the false negative rate (FNR) in our DNM detection pipeline, we generated simulated data based on callable genome regions of three selected offspring. For each of the three individuals, we made an artificial genome based on the callable genome regions of that individual and put a specific number of simulated DNMs within the artificial genome. We next generated 80× simulated paired-end 150 bp reads for each selected individual using ART (Huang, et al. 2012) and re-ran the full alignment and DNM detection workflow. The detected DNMs were then compared with the known positions of simulated DNMs to estimate the FNR (the equation below) for each selected offspring independently. The final FNR value was reported as the average across the three simulations (details given in **Supplementary Table 11**).

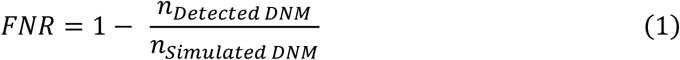

### *De novo* mutation rate estimation

The per-site-per-generation mutation rate (µ) was calculated as:

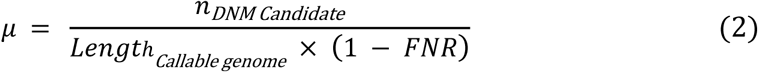

The callable genome size (Length_Callable genome_) was independently determined for each offspring using the procedures described above. The FNR of the DNM detection pipeline was estimated from the simulation described above. All candidate DNMs were visually reviewed in IGV (v2.19.2) (Thorvaldsdottir, et al. 2013), and a subset was further validated by Amplicon-seq. These quality-control steps ensured high confidence in DNM identification. As all putative false positive DNMs were removed, we did not apply false discovery rate (FDR) correction for mutation rate estimates.

### Analysis of nucleotide diversity in amphioxus populations

To estimate nucleotide diversity in *B. floridae* populations, we re-analyzed previously published population data. First, we downloaded whole-genome sequencing data of 35 *B. floridae* individuals (Huang, et al. 2023) from NCBI’s SRA database. The short-read alignment and variant calling largely followed the method of Huang et al. (Huang, et al. 2023), with key adaptations. Specifically, the *B. floridae* genome assembly of that study (Huang, et al. 2023) was utilized as the reference, with reads mapped using BWA-MEM2 (v2.2.1) (Vasimuddin, et al. 2019) under default parameters optimized for vertebrate genomes. We then generated gVCF files for each individual using GATK4 (v4.6.1.0) (Van der Auwera and O’Connor 2020) HaplotypeCaller, setting the “-ERC BP_RESOLUTION” to include non-variant sites in the output, and adjusting parameters to reflect the heterozygosity of amphioxus by parameters “--heterozygosity to 0.03 and --indel-heterozygosity to 0.003”. The sequence alignment and variant detection workflows were integrated into an automated nf-core/sarek pipeline (v3.4.0) (Garcia, et al. 2020) with key parameters fine-tuned, thereby maximizing the utilization of parallel computing resources.

For variant sites, we performed initial filtering using GATK4 VariantFiltration on SNPs and INDELs independently, excluding SNPs with QUAL<30, QD<2, FS>60, MQ<40, MQRankSum<-12.5, Read PosRankSum<-8 or SOR>3 and excluding INDELs with QUAL<30, QD<2, FS>200, ReadPosRankSum<-20, and SOR>10 (Mirchandani, et al. 2024). For invariant sites, we filtered based on quality (“QUAL>30”) and the fraction of missing genotypes at a site (“F_MISSING<0.25”).

To estimate nucleotide diversity at synonymous sites (π_S_), we used the script dNdSpiNpiS (v1.0) (https://kimura.univ-montp2.fr/PopPhyl/index.php?section=tools) with the default parameters. To extract phase information for the CDS features, we used GenomeTools (gt gff3 -sort -tidy -retainids). Based on the VCF file, we then generated the diploid fasta sequence for each individual using the FastaAlternateReferenceMaker module from GATK (v4.6.1.0) (Van der Auwera and O’Connor 2020). We then intersected these fasta sequences with the gene annotation GFF file to keep only CDS with SNPs. For each CDS, individuals with missing data were excluded when estimating π_S_. As for *π*, these estimates were corrected to account for truly invariant sites versus those resulting from insufficient sequencing depth. We used the invariant sites generated above to estimate the expected percentage of bases called per CDS and applied this measure as a correction factor for the π_S_ estimates. The mean diversities π_S_ of the CDS were then calculated for the whole genome.

### Analysis of the effective population size

The average long-term *N_e_* of *B. floridae* was estimated using the formula *N_e_* = *π*/(4*μ*) (Kimura 1983), where *μ* was the mutation rate estimated by our work and π_S_ was estimated with data from a previous study (Huang, et al. 2023) (**Supplementary Table 8**). For comparison, *μ* and *π* data of vertebrates were from Bergeron et al. (Bergeron, et al. 2023).

### Aligning parental genome assemblies to the published *B. floridae* genome

Owing to inherent limitations of short-read sequencing, the assembly of parental genomes consisted of primarily short contigs. Defining orthologous relationships between contigs of two parents in a specific locus based solely on pairwise alignments of contigs, as previously mentioned, can be complicated by the variability in the lengths of assembled contigs derived from that locus. To address these issues, we aligned the contigs of parental genomes to a published *B. floridae* haploid reference genome (Huang, et al. 2023). After testing multiple alignment tools under various parameter configurations, Minimap2 (v2.24) (Li 2021) was selected as the optimal aligner, with empirically optimized parameters: “-x asm20 --secondary=no”. Resultant alignments were processed through sequential refinement: (1) coordinate sorting via SAMtools (v1.21) (Li, et al. 2009) and (2) removal of multi-mapping regions using Sambamba (v0.8.2) (Tarasov, et al. 2015). A custom script was employed to establish the mapping relationship between each parental genome and the haploid reference genome. This approach yielded a multiple genome alignment that aligned positions across the paternal, maternal, and reference genomes. As a result, the majority of parental genomic positions were successfully anchored onto the reference, thereby obtaining orthologous genomic coordinates across the parental genomes and facilitating subsequent analyses dependent on chromosomal coordinates.

### Comparative analysis of mutation rates in different contexts

To compare mutation rates in different contexts (e.g., different annotation groups), we utilized the coordinate mapping relationships described in the previous section. We mapped the gene functional annotations from the published *B. floridae* reference genome (Huang, et al. 2023) onto the parental genomes. In doing so, the coordinates for exonic, intronic, and intergenic regions were converted to those of the parental genomes. This allowed us to determine the lengths of each gene feature where DNMs and callable genome regions were present, enabling the calculation of annotation-specific mutation rates for each individual.

To generate the mutation spectrum, the DNMs were also stratified into six groups according to their mutation types (A>C/T>G, A>T/T>A, A>G/T>C, C>A/G>T, C>G/G>C and C>T/G>A), with a special group for C>T mutations at CpG sites. We extracted CpG positions from the parental genomes, thereby allowing us to distinguish between CpG and non-CpG sites.

### Identifying the candidate PZMs

Raw VCF files of parent-offspring trios were obtained from the section “Alignment of offspring and parental reads to the parental reference genome and variant calling”. PZMs were retained only if they met all the following criteria: (1) the variant was located within the predefined callable genome regions; (2) the offspring harbored two alleles for a specific site in a parental contig with a total read depth > 10; (3) there were at most two alleles at the variant site in each parental BAM file; (4) the variant with no proximity to indels (outside ±20 bp of indels); (5) the variant was not located in the regions experiencing putative gene conversions; (6) excluding candidate PZMs where the derived allele had ≥2 reads in any other individual.

For each candidate PZM, IGV screenshots were generated for both offspring and parents using the method outlined in the “Identifying the candidate DNMs” section. Subsequent manual inspection led to the removal of any candidate PZMs if they met one or more of the following criteria (1) structural variant/recombination breakpoint neighborhoods; (2) variants were found to arise from misassigned parental origins of contigs, upon manual review.

### Amplicon-seq validation

To assess the reliability of DNMs across different data sources, we randomly selected a subset (**Supplementary Table 7**) for validation by amplifying target regions with specific primers, followed by amplicon sequencing for genotyping. The validation workflow comprised three main steps: primer design, *in silico* PCR, and multiplex PCR amplification coupled with amplicon sequencing.

In the primer design phase, due to the high sequence polymorphism observed in *B. floridae* (∼4%), it was challenging to simultaneously amplify parental alleles using a single pair of primers targeting conserved regions; therefore, it became necessary to design allele-specific primers for each allele from the two parents. Specifically, we adopted the framework of the NGS-PrimerPlex (v1.3.4) (Kechin, et al. 2020) multiplex PCR primer design process, with modifications tailored to our specific requirements. Since we needed to amplify all relevant alleles independently in offspring and parents, we chose the parental reference genome as the reference for primer design to maximize the number of alleles amplified by a single primer pair. For each locus, we used the mapping relationships between parental genomes, as described in the section “Aligning parental genome assemblies to the published *B. floridae* genome”. This allowed us to extract the target alleles harboring the DNMs and the orthologous alleles from the other parent. To avoid software misinterpretation of homologous regions as multiple alignments, we removed non-target contigs from the reference genome. We then used the processed parental reference genome for each individual together with the DNM positions to run NGS-PrimerPlex (parameters given in the **Supplementary Table 5**). After executing the pipeline, we selected the top five multiplex PCR primer sets with the highest primer scores as candidate primer combinations for the subsequent steps (**Supplementary Table 6**).

*In silico* PCR was employed to confirm that primers designed for multiplex amplification can effectively amplify all parental alleles at the loci harboring selected DNMs, using *in_silico_pcr.pl* (v0.6, https://github.com/egonozer/in_silico_pcr). To optimize efficiency, only the contig harboring the DNM and orthologous contigs were extracted as the reference genome for *in silico* PCR. If *in silico* PCR showed incomplete allele amplification, we designed additional primers for the unamplified contig. These primers were added to the multiplex primer set, and the process was repeated until all target fragments could be amplified.

For multiplex PCR, we used the gDNA templates of offsprings and parents stored previously and performed PCR amplification with a multiplex PCR kit (VAZYME, CN: PM101-02) optimized for both sensitivity and specificity; the resulting products are then subjected to agarose gel electrophoresis, and finally, DNA fragments ranging from 150 to 200 bp were excised from agarose gel following electrophoresis and subsequently purified using gel extraction kit. Library quality was assessed by fragment analysis using the Qsep-400 and quantified for concentration using the Qubit 3.0 Fluorometer (Thermo Fisher Scientific, Waltham, MA). Library construction was performed using the VAHTS^®^ Universal Plus DNA Library Prep Kit for Illumina ND617, following the manufacturer’s protocol. Libraries meeting the quality control criteria were subsequently sequenced on the Illumina NovaSeq 6000 platform.

## Data Availability

The raw sequence data reported in this paper have been deposited in the Genome Sequence Archive (GSA: CRA027444) that are publicly accessible at https://ngdc.cncb.ac.cn/gsa. The raw sequence data of previously sequenced populations can be accessed under the BioProject with accession number: PRJNA602496. Genome assemblies produced by Platanus-allee and the amplicon sequencing data have been deposited in Science Data Bank (ScienceDB) repository (https://doi.org/10.57760/sciencedb.27648). Custom scripts used are available online at https://github.com/xuejinghub/Amphioxus_denovo_mutation_rate.

## Author contributions

**Jing Xue**: Investigation, Methodology, Formal analysis, Writing - Original draft, Writing - Review & Editing. **Lei Tao**: Investigation, Methodology, Formal analysis, Writing - Review & Editing. **Junwei Cao**: Data Curation, Investigation. **Guang Li**: Conceptualization, Data Curation, Resources, Writing - Review & Editing, Supervision. **Cai Li**: Conceptualization, Resources, Methodology, Writing - Review & Editing, Supervision.

## Acknowledgments

We thank Dr. Luohao Xu for his assistance in obtaining previous genomic data of amphioxus populations. We thank Dr. Chung-I Wu and Dr. Luohao Xu for their comments on the manuscript. This work was supported by National Natural Science Foundation of China (32470690), Natural Science Foundation of Fujian Province of China (2022J06004), State Key Laboratory of Biocontrol and Guangdong Provincial Key Laboratory for Aquatic Economic Animals.

## Supplementary Tables and Figures

Supplementary Tables are provided in a separate Excel file, and Supplementary Figures are shown below.

**Supplementary Figure 1.**
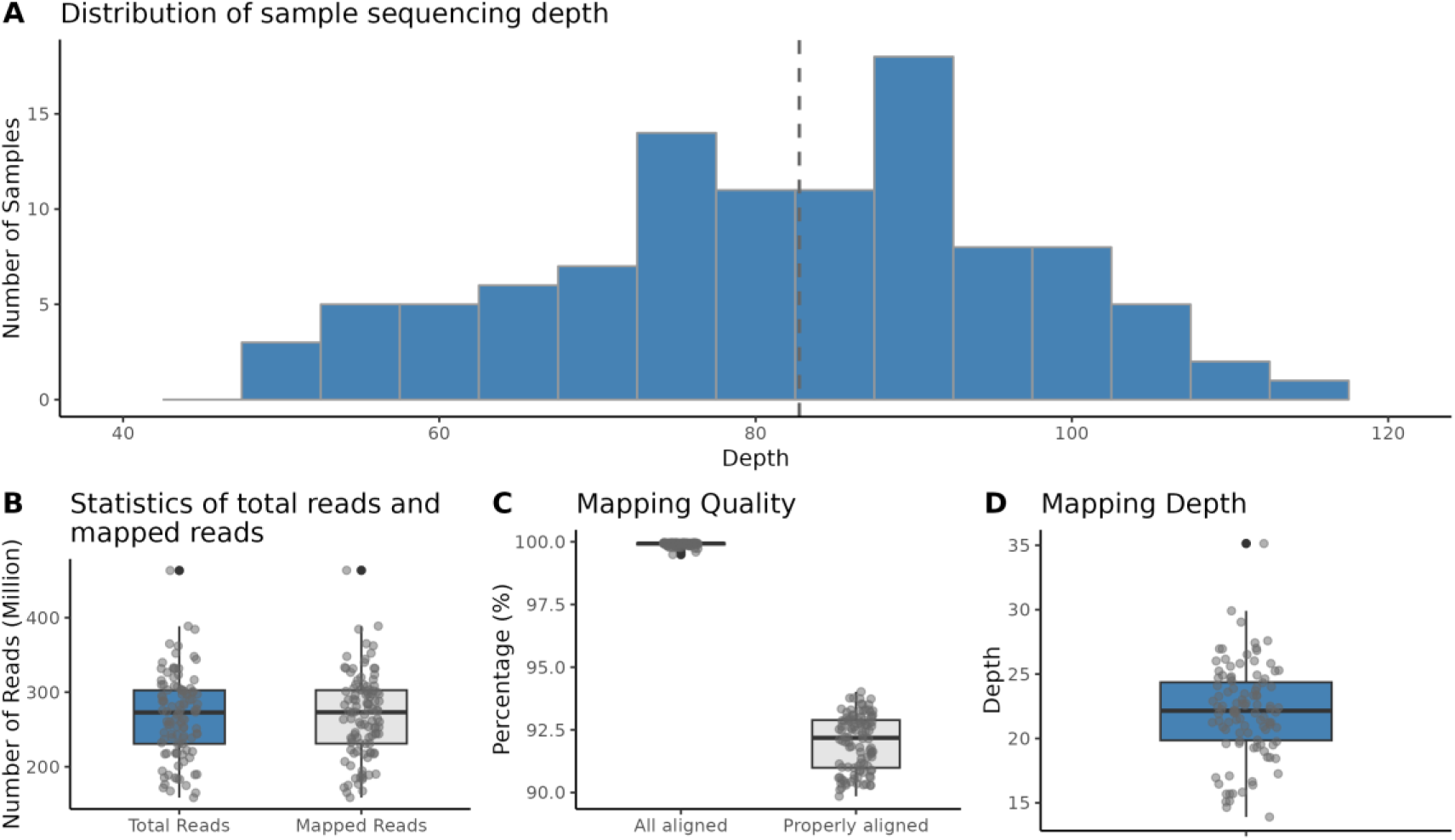
Statistics of sequencing and mapping results. **(A)** Distribution of sequencing depths of all samples. (**B**) Boxplots showing the numbers of all reads and the numbers of mapped reads against the parental genomes. (**C**) Boxplots showing the percentages of all reads that were mapped to the parental genome, and the percentage of properly aligned reads. ‘Properly aligned’ means that paired-end reads were aligned to the same sequence with an insert size of ≤ 1,000 bp. (**D**) Boxplot showing average read depths of samples based on the alignments against the parental genome.

**Supplementary Figure 2.**
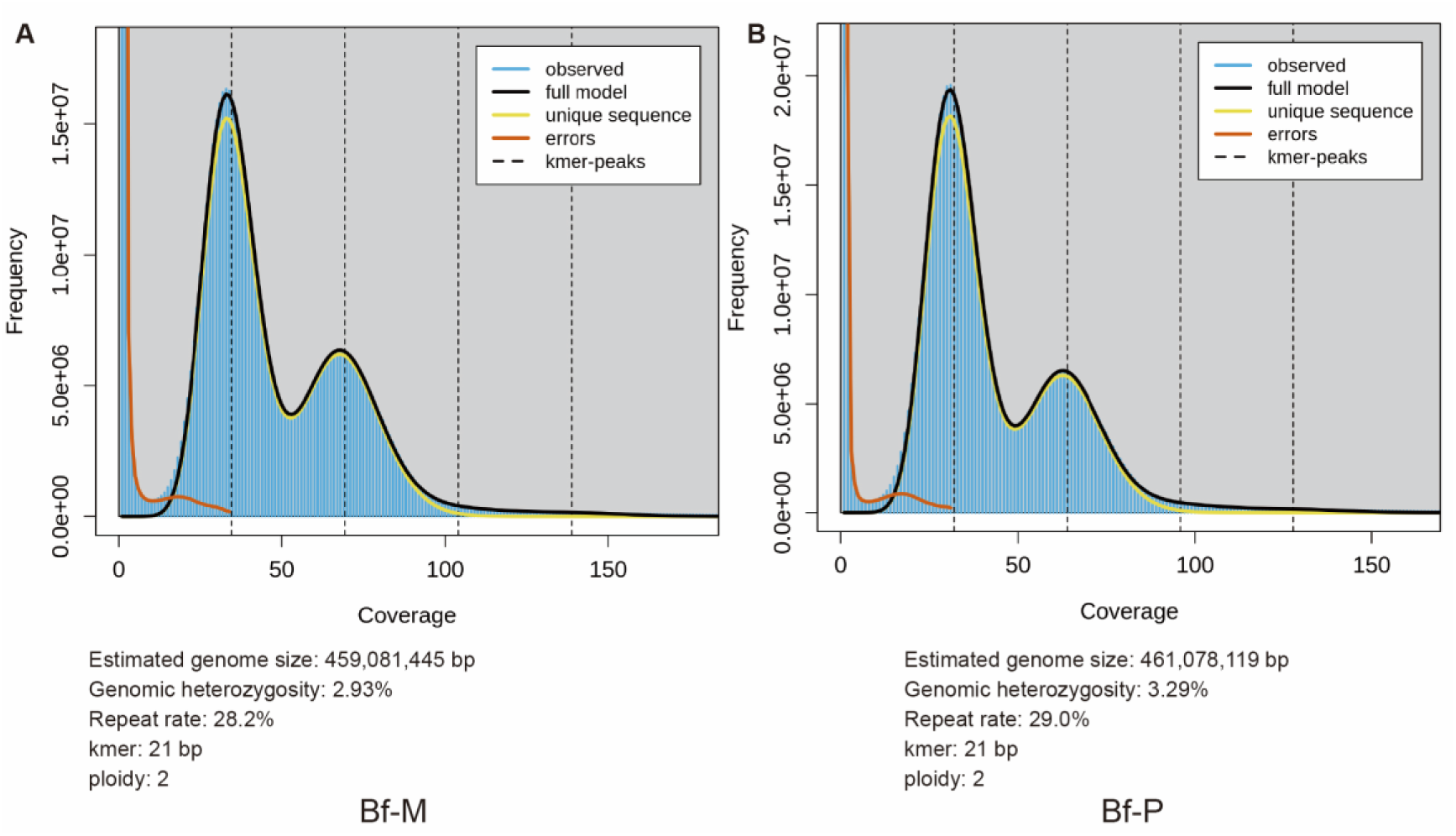
Genome characteristics of *B. floridae* parental genomes based on 21-mer analysis. (**A**) Maternal genome. (**B**) Paternal genome. The bottom of each panel displays genome characteristics inferred from 21-mer analysis, including estimated genome size, genomic heterozygosity, repeat rate, *k*-mer size and ploidy.

**Supplementary Figure 3.**
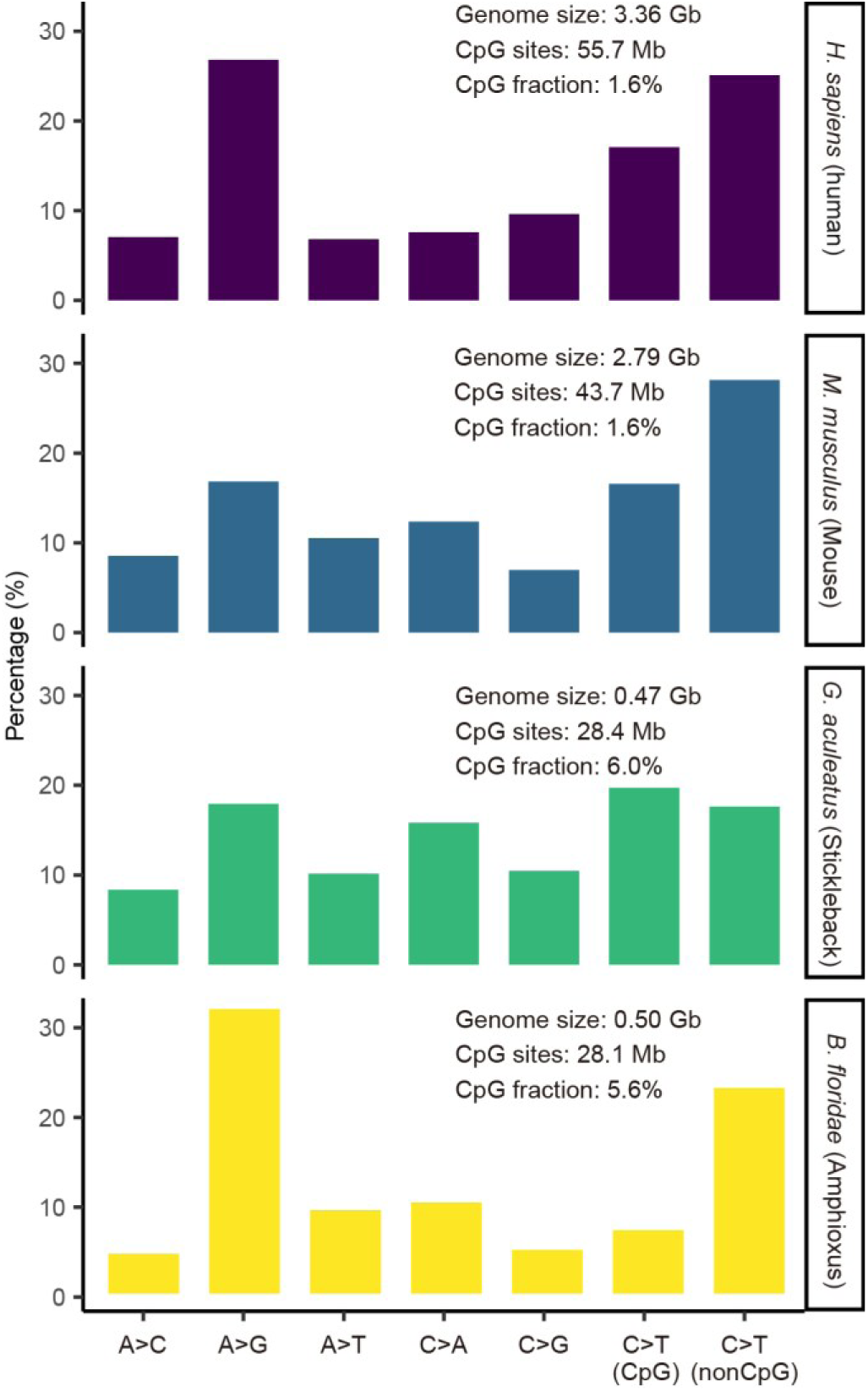
Mutation spectra of *B. floridae* and representative vertebrate species. *H. sapiens* (Human), purple; *M. musculus* (Mouse), blue; *G. aculeatus* (Stickleback), green; *B. floridae* (Amphioxus), yellow.

**Supplementary Figure 4.**
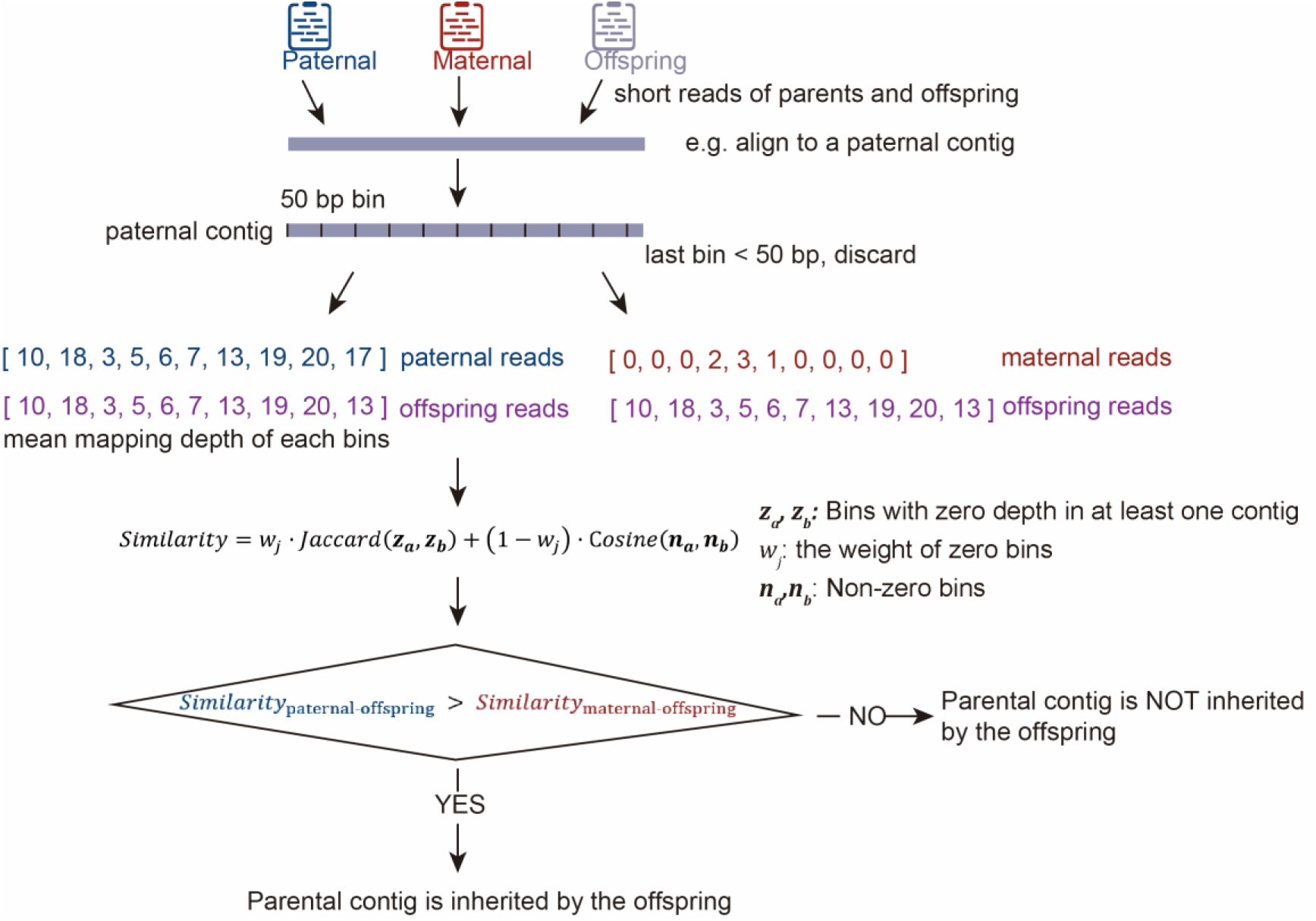
Inferring inheritance states of parental contigs in offspring. First, we aligned parents’ and offsprings’ WGS reads to the parental reference genome. In the figure, we chose a paternal contig as an example. To quantify depth distribution, the paternal contig was divided into 50-bp bins, discarding any terminal bin shorter than 50 bp. The mean depth within each bin was calculated for the offspring, the mother and the father, respectively, generating a depth distribution vector per individual across the paternal contig. The similarity between offspring and parental depth vectors was quantified using a weighted measure combining: (1) Cosine similarity for bins with non-zero depths in both individuals being compared, and (2) Jaccard similarity for bins with zero coverage in at least one of the two individuals (accounting for regions with no aligned reads). The paternal contig was considered inherited by an offspring if the offspring’s depth distribution vector showed greater similarity to that of the father than to the mother.

**Supplementary Figure 5.**
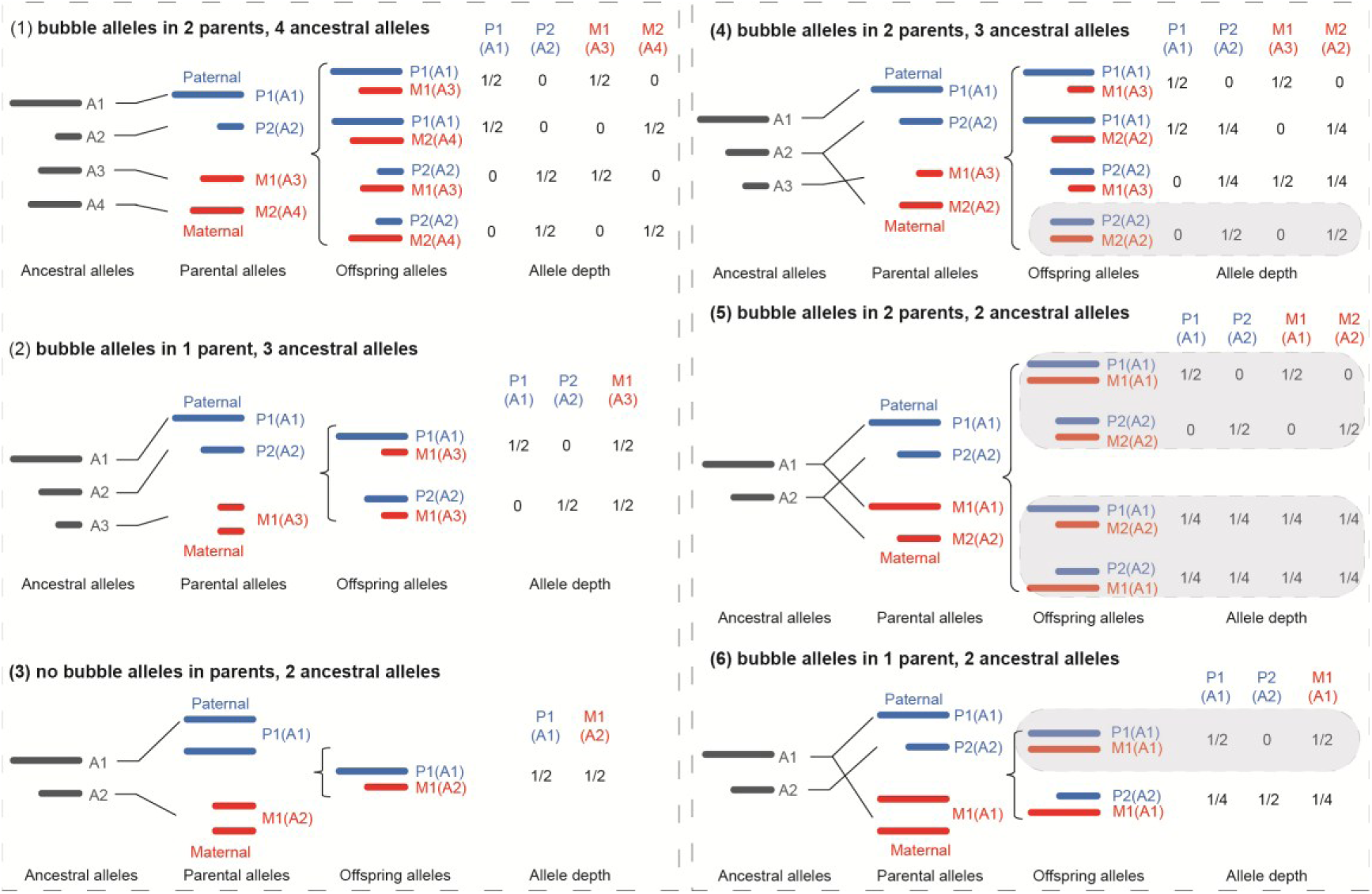
Possible allele combinations in the paternal and maternal genomes at a locus and possible resulting read depths of alleles in the offspring. Distinct alleles (assembled contigs) are named A1, A2, etc. P1, P2, M1 and M2 are names of paternal and maternal alleles. The value of an allele depth represents the relative ratio of read depth of the specific allele to an offspring’s average read depth. For scenarios (1) - (3), the two parents do not have shared (highly similar) alleles, the inheritance states of alleles can be determined and the offspring alleles can be phased. In contrast, for scenarios (4) - (6), when some alleles that are highly similar between parents are passed to an offspring (see shaded areas in the figure), the inheritance states of alleles cannot be reliably determined or offspring alleles cannot be phased. Therefore, the genomic regions in the offspring matching the conditions in shaded areas were not included in the callable genome.

